# Interruption of thymic activity in adults improves responses to tumor immunotherapy

**DOI:** 10.1101/2020.01.08.899484

**Authors:** José Almeida-Santos, Marie-Louise Bergman, Inês Amendoeira Cabral, Jocelyne Demengeot

**Affiliations:** Instituto Gulbenkian de Ciência, Rua da quinta grande 6, 2780 Oeiras, Portugal

## Abstract

The thymus produces precursors of both effectors and regulatory T cells (Tconv and Treg, respectively) whose interactions prevents autoimmunity while allowing efficient protective immune responses. Tumors express a composite of self- and tumor-specific antigens and engage both Tconv and Treg cells. Along the aging process, the thymus involutes, and tumor incidence increases, a correlation proposed previously to be causal and the result of effector cell decline. In this work, we directly tested whether interruption of thymic activity in adult mice affects Foxp3 expressing Treg composition and function, and alters tumor immune surveillance. Young adult mice, on two different genetic backgrounds, were surgically thymectomized (TxT) and analyzed or challenged 2 months later. Cellular analysis revealed a 10-fold decrease in both Tconv and Treg numbers and a bias for activated cells. The persisting Treg displayed reduced stability of Foxp3 expression and, as a population, showed compromised return to homeostasis upon induced perturbations. We next tested the growth of three tumor models from different origin and presenting distinct degrees of spontaneous immunogenicity. In none of these conditions adult TxT facilitated tumor growth. Rather TxT enhanced the efficacy of anti-tumor immunotherapies targeting Treg and/or the checkpoint CTLA4, as evidenced by increased frequency of responder mice and decreased intra-tumoral Treg to CD8^+^IFNγ^+^ cell ratio. Together, our findings point to a scenario where abrogation of thymic activities affects preferentially the regulatory over the ridding arm of the immune activities elicited by tumors, and argues that higher incidence of tumors with age cannot be solely attributed to thymic output decline.

## Introduction

The thymus exports T cells that colonize peripheral lymphoid and non-lymphoid organs and continuously replace dying T cells, ensuring the maintenance of a diverse reactive repertoire. Thymic atrophy in response to various biological stresses, and involution with age, leads to alteration in peripheral T cell composition. T cells can be subdivided in two broad types, those “conventional” or effector cells, participating in the ridding functions of the immune system, and those endowed with regulatory functions, which maintain at check effector cells, and most notably prevent deleterious autoimmunity. The major subset of regulatory T cells (Treg) expresses the master transcription factor Foxp3, whose mutation leads to severe, lethal multiorgan autoimmunity in mice and humans (1-3). Several experimental and theoretical works have attempted to formalize which mechanisms ensure the adequate balance between regulatory and effector T cells number and composition in health. Overall the two cell subsets share only few TCR sequences when analyzed at the nucleotide level (4), and the most consensual model remains that they must be docking on the same antigen presenting cells for regulation to occur and be reinforced (5).

Most analyses on the impact on health of thymic surgical ablation, physiological atrophy or involution, focused on the ensuing restricted repertoire of effector T cells, which satisfyingly explains a reduced efficiency in targeting infectious agents. However, thymic output is also required to replenish the pool of peripheral Treg, through both the production of Treg already differentiated in the thymus (tTreg) (2) and the production of immature T cells that are the preferential precursors of *bona fide* Treg differentiated in the periphery (6-8).

Cancer immune surveillance involves both the regulatory and effector arms of the immune system (9). Cancer cells are derived from tissues that in health are not targeted by the ridding function of the immune system as these are maintained at check by Treg, among other suppressor mechanisms. In turn, beneficial anti-tumor immune responses have similarities with autoimmunity, as illustrated by the emergence of autoimmune manifestations upon immunotherapies (10, 11). More recent advances in genome analysis has revealed that most tumors express neo-antigens, encoded by mutated genes. Tumors bearing mutator mutations accumulate multiple neo-antigens, and appear the best responders to immune therapies (12). Nevertheless, Treg interacting with tumor antigens are identifiable and these derive both from tTreg and pTreg (13, 14).

In a recent publication, Palmer et al (15) argued that the increased incidence of cancer with age is not caused by gradual accumulation of mutations as commonly accepted, but rather by a decline in the immune system, that the authors attribute to a decline in thymic T cell output. In their mathematical model, all T cells are treated as effector cells. Given the evidence that the thymus produces precursors of both effectors and Treg, and that the outcome of an immune response is the result of effector and Treg interactions, predicting the impact of thymic output interruption on tumor immune surveillance is non-trivial.

In the present work, we directly tested whether interruption of thymic activity in adult mice affects Treg composition and function in the periphery. Mice were surgically thymectomized (TxT) between 5 and 6 weeks of age to abrogate thymic output, and analyzed 2 months later. We first document that as for conventional T cells, Treg numbers decrease and the remaining cell population is biased for activated Treg. The persisting Treg are less stable and respond worse to induced homeostatic perturbations. We further tested the impact of adult TxT on the growth of three tumor models in two mouse strains of different genetic backgrounds. In none of the circumstances TxT facilitated tumor growth, and in some instances, TxT enhanced the efficacy of anti-tumor immunotherapies.

## Results

### Adult thymectomy alters both conventional and regulatory T cell composition

We first analyzed the T cell composition in C57Bl/6 (B6) animals that had been submitted to adult thymectomy (TxT) or sham surgery (Sham) 2 months earlier (**Fig. 1A**). As expected, recent thymic emigrants identified as Qa-2^low^ were drastically decreased in TxT mice (**Fig. S1A**). Naïve CD4 and CD8 cells, as well as central memory Treg (cTreg) are identified as CD44^-^CD62L^+^, while activated or effector memory CD4, CD8 and Treg (eTreg) cells are identified as CD44^+^CD62L^-^ (16). Assessing both the frequency and absolute cell numbers revealed that the log decrease in cell numbers observed in TxT mice is mostly imputable to naïve T and cTreg cells, while the number of activated T and eTreg cells remains unaltered (**Fig. 1B, C)**. Of note, when monitoring the cellular decline upon TxT in BALB/c (Ba) mice, the decrease in naïve CD4 or CD8 T cells and in cTreg numbers was much less marked and delayed, when compared to B6 animals (**Fig. S1B**). The extent of naïve T cell decline in B6 TxT mice was in the same range as in 15-month-old animals (**Fig S1C**). TxT and Sham animals displayed similar frequency of Ki67+ cells in each T cell subset **(Fig. S1D)**, confirming peripheral proliferation does not compensate for cessation of thymic output in mice (17). The frequency of activated cells that produced interferon-γ (IFN-γ) or tumor necrosis factor α (TNF-α) were not decreased in TxT mice when compared to Sham animals **(Fig. 1D**). We further sub-phenotyped Treg according to various markers (**Fig. 1E**). Expression of Helios and Nrp1 has been proposed, and debated, as markers of Treg differentiated in the thymus (tTreg) *versus* in the periphery (pTreg) (18). The frequency of cTreg expressing Nrp1 was increased in TxT mice, possibly indicating a lack of Nrp1^-^ pTreg, while Helios^+^ cells were underrepresented. The intensity of Foxp3 expression in cTreg was slightly but significantly reduced in TxT animals, suggesting less robust stability or function. Analysis of other markers associated with Treg function, such as CD25, CTLA4 and TIGIT, revealed no significant differences between TxT and control mice, either for cTreg or eTreg. Together, these data indicate that interruption of thymic activity affects Treg composition in the periphery.

**Figure 1.**
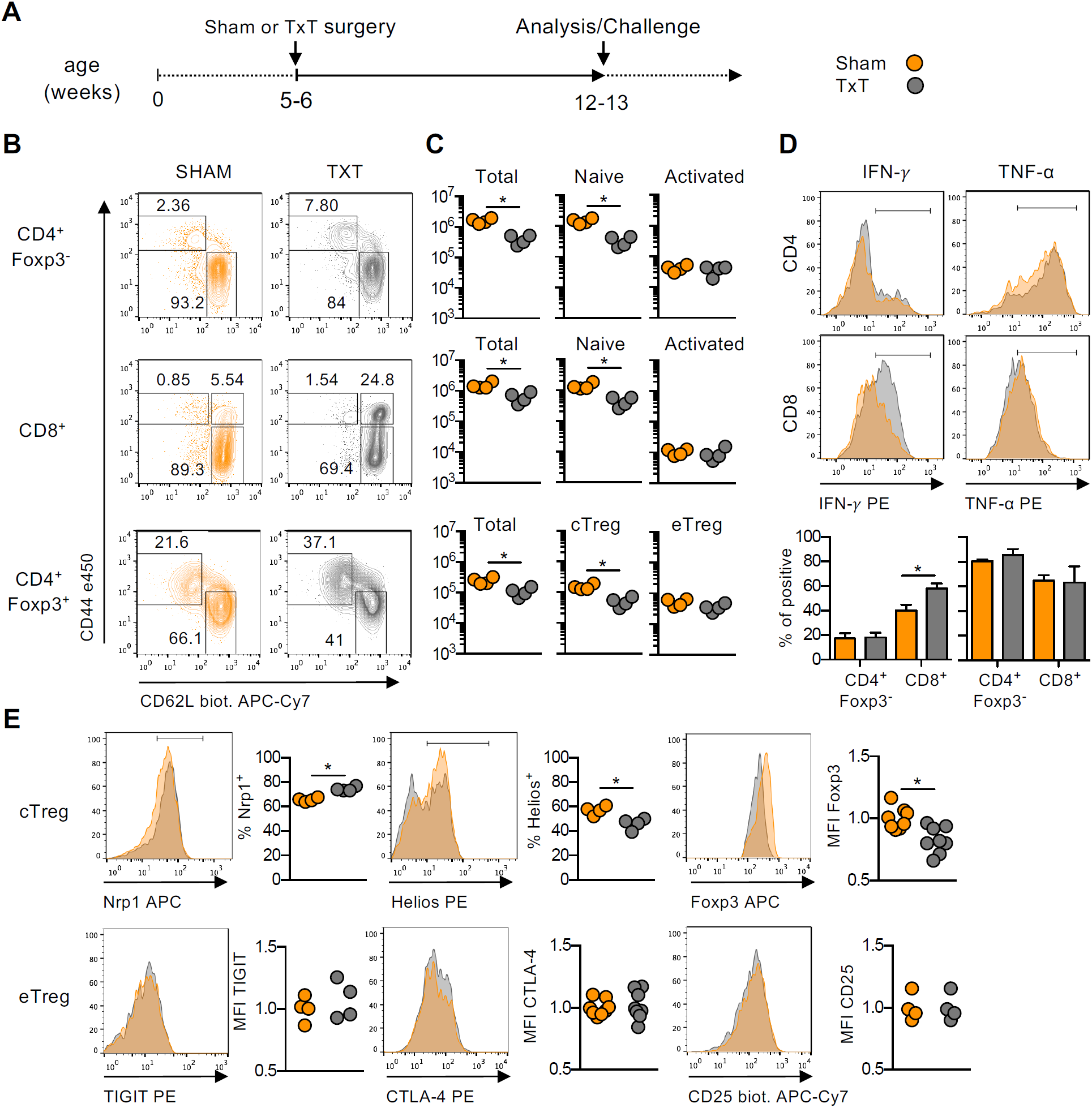
Adult thymectomy alters both conventional and regulatory T cell composition. **A**) Schematic representation of thymectomized (TxT) and sham-operated (Sham) mice. **B-D**) Analysis of inguinal lymph nodes (LN) for expression profile of CD44 and CD62L inside Foxp3^-^CD4^+^, Foxp3^-^CD8^+^ and Foxp3^+^CD4^+^ cells (B), total numbers of T cells belonging to each subset (C) and interferon-*γ*(IFN-*γ*) and TNF-α cytokine production in CD44^+^ CD4^+^ and CD8^+^ cells from Sham and TxT mice (D). Representative of 2 independent experiments (n=3-4). **E**) phenotypic characterization of cTreg (Foxp3^+^ CD44^-^ CD62L^+^) and eTreg (Foxp3^+^ CD44^+^ CD62L^-^) from Sham and TxT mice, representative of 2 independent experiments (n=3-4), except for Foxp3 and CTLA4, pool of 2 independent experiments (n=3-4). MFI: mean fluorescence intensity (normalized to average Sham values). Statistical analysis performed using nonparametric T-test *P < 0.05.

### Functional alteration of Treg upon adult thymectomy

We next tested the function and stability of Treg isolated from TxT or Sham mice by adoptive transfer. Conventional CD4 and CD8 cells were purified from unmanipulated donors and injected alone or together with Treg in T cell deficient animals (**Fig. 2A**). In such adoptive transfer experiments conventional T cells expand vigorously and acquire an activated phenotype if injected alone, a process inhibited by the co-transfer of Treg. Treg isolated from TxT mice performed similarly to, or worse than Treg isolated from Sham mice, in controlling CD4 or CD8 cells, respectively. Adoptive transfer of Treg cells alone serves as an assay to test their plasticity/stability, measured through the frequency of cells that lose Foxp3 expression along the experiment. Tregs isolated from TxT mice, when compared to Sham donors, were less stable in this assay. Loss of Foxp3 expression in both types of Treg was equally prevented by co-transfer of CD4 cells, a mechanism shown before to involve IL-2 secreted by effector cells (19), and not by co-transfer of CD8 cells. Adoptive transfer of conventional CD4 T cells alone serves also as an assay to measure differentiation of Treg in the periphery. Our previous work demonstrated that naïve cells isolated from TxT mice failed to express Foxp3 upon adoptive transfer into lympho-deficient mice (7). We extended this finding by performing adoptive transfer into lymphoreplete animals. While donor cells and newly differentiated Treg are barely detectable in lymphoreplete recipients, these are readily measurable when using DEREG recipient animals partially deleted of endogenous Treg (8). In the latter assay, as in lympho-deficient mice, cells isolated from TxT mice performed poorly **(Fig 2B)**. Taken together, these data indicate that TxT mice bear compromised Treg that cannot be efficiently renewed through peripheral differentiation.

**Figure 2.**
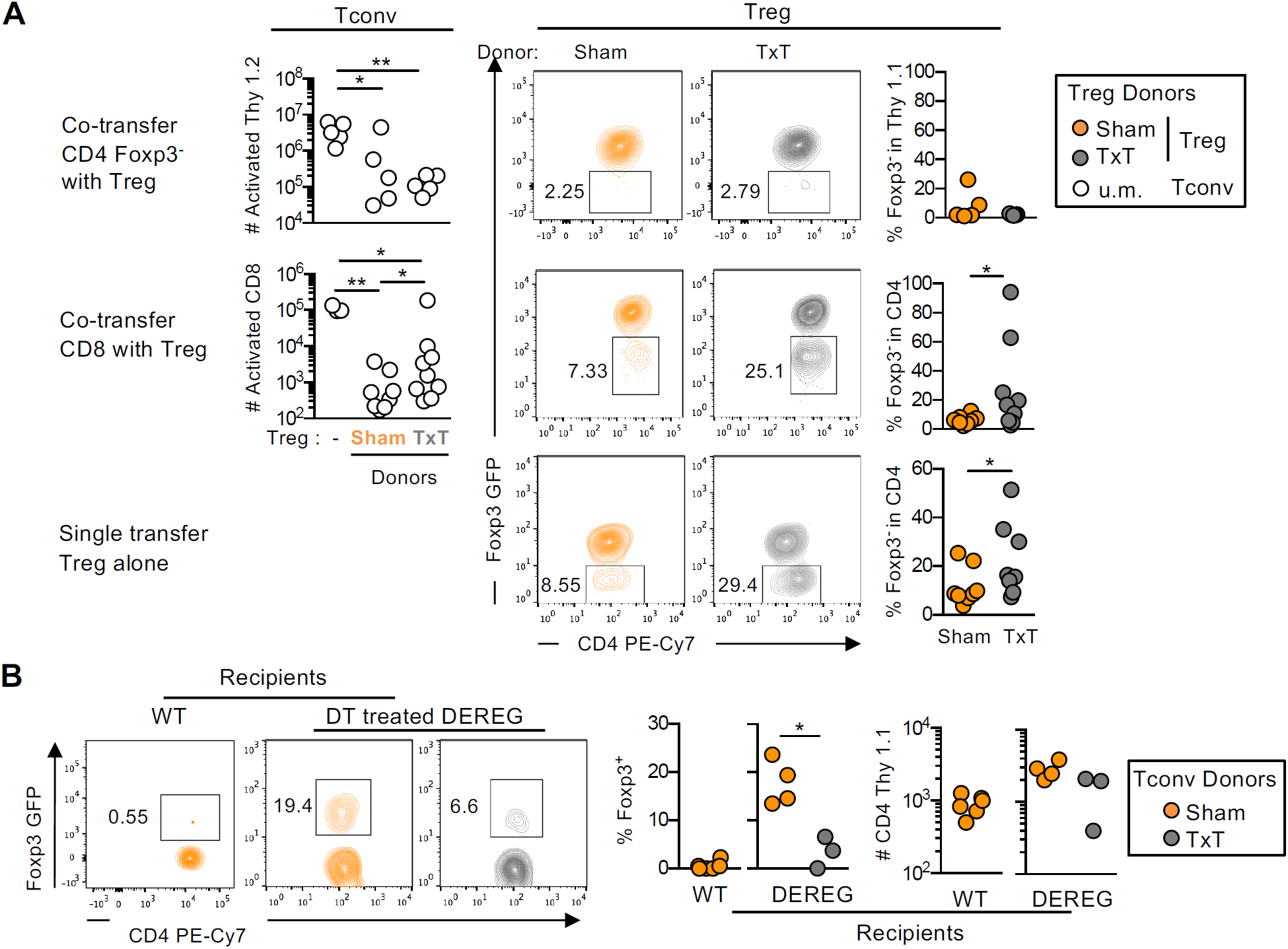
Functional alteration of Treg upon adult thymectomy. **A**) Regulation and stability. Sham or TxT Foxp3^GFP^ mice were donors of CD4^+^ Foxp3^+^ Treg, euthymic unmanipulated (u.m.) mice were donors of CD4^+^Foxp3^-^ and CD8^+^ conventional T cells (Tconv). TCRβ^-/-^ recipients received either Thy1.2 CD4^+^Foxp3^-^ T cells as target cells transferred alone or together with Thy1.1 Treg (upper panels), CD8 T cells as target transferred alone or together with Treg (middle), or a single transfer of Treg (bottom). Shown in the left panels (Tconv) are the numbers of activated Thy 1.2 CD4 cells (upper), or CD8+ T cells (middle) recovered 1 month after adoptive transfer. Shown in the right panels are the analysis of originally Treg in each adoptive transfer: representative flow cytometry analysis and frequency of Treg that lost Foxp3 expression. Upper panel one experiment (n= 5), middle and lower panels pool of 2 independent experiments (each n=4-5). **B**) De novo differentiation. Single transfer of CD4 naïve cells isolated from Sham or TxT Thy 1.1 Foxp3^GFP^ donors into Thy 1.2 wild type (WT) or diphtheria toxin (DT) treated DEREG recipients. DT was administrated twice, the day before and the day of adoptive transfer. Shown is a representative flow cytometry analysis and the percentage of Foxp3^+^ cells recovered 1 month after adoptive transfer, representative of 2 independent experiments (n=3-4). Statistical analysis was performed using nonparametric Mann Whitney T-test. *P < 0.05, **P < 0.005

### Altered return to homeostasis upon Treg depletion in thymectomized mice

DEREG mice, which carry a BAC transgene covering the foxp3 locus knocked-in to encode DTR and GFP, offer a practical model to study Treg dynamics upon perturbation. Punctual injection of diphtheria toxin (DT), depletes all GFP^+^ Treg in lymphoid and non-lymphoid tissues but spare DT resistant GFP^-^ Treg that are Foxp3^+^ but do not express the BAC (**Fig. S2A**). The rebound of Treg is rather fast and biphasic, with the rapid emergence (presumably expansion) of GFP-Treg followed by a slower reappearance (presumably *de novo* differentiation) of GFP^+^ Treg (**Fig. S2B**).

In this system GFP^-^ Treg have been shown to be rather CD25^low^, highly cycling, hyperactivated and exhausted (20). By crossing DEREG mice with Foxp3^hCD2^ animals bearing different congenic markers (CD45.2 and CD45.1, respectively), we produced a tool that allows functional analysis of GFP^-^ Treg. Probing this system in adoptive transfer experiments revealed that hCD2^+^GFP^-^ cells transferred into DT treated DEREG animals did not acquire GFP expression in a time frame during which endogenous GFP^+^ cells had already emerged **(Fig. S2C**). We also evidenced that the majority of Treg differentiated in the periphery upon adoptive transfer of Foxp3^-^ cells were GFP expressors (**Fig. S2D**). This analysis confirms that GFP^-^ Treg in DEREG mice are not precursors of GFP^+^ *bona fide* Treg, and are likely to represent aged Treg.

Monitoring Treg rebound in the blood of TxT and Sham DEREG over 25 days after DT treatment (**Fig 3A**) revealed similar kinetics when testing the frequency and number of total Treg. However, in TxT animals GFP^+^ cells were underrepresented, and early rebounding Treg expressed less CD25. Lymph node analysis at day 6 post DT injection confirmed the disbalanced rebound of Treg in TxT compared to Sham mice (**Fig 3B**). Finally, analysis 35 days post DT treatment (**Fig 3C**) evidenced that while Sham animals return to homeostasis replenishing both cTreg and eTreg compartments, TxT animals contained mostly eTreg. Moreover, eTreg in TxT animals appear overactivated with elevated levels of CTLA4 expression. All together these results indicate that induced perturbation of Treg has stronger cellular impact in TxT mice.

**Figure 3.**
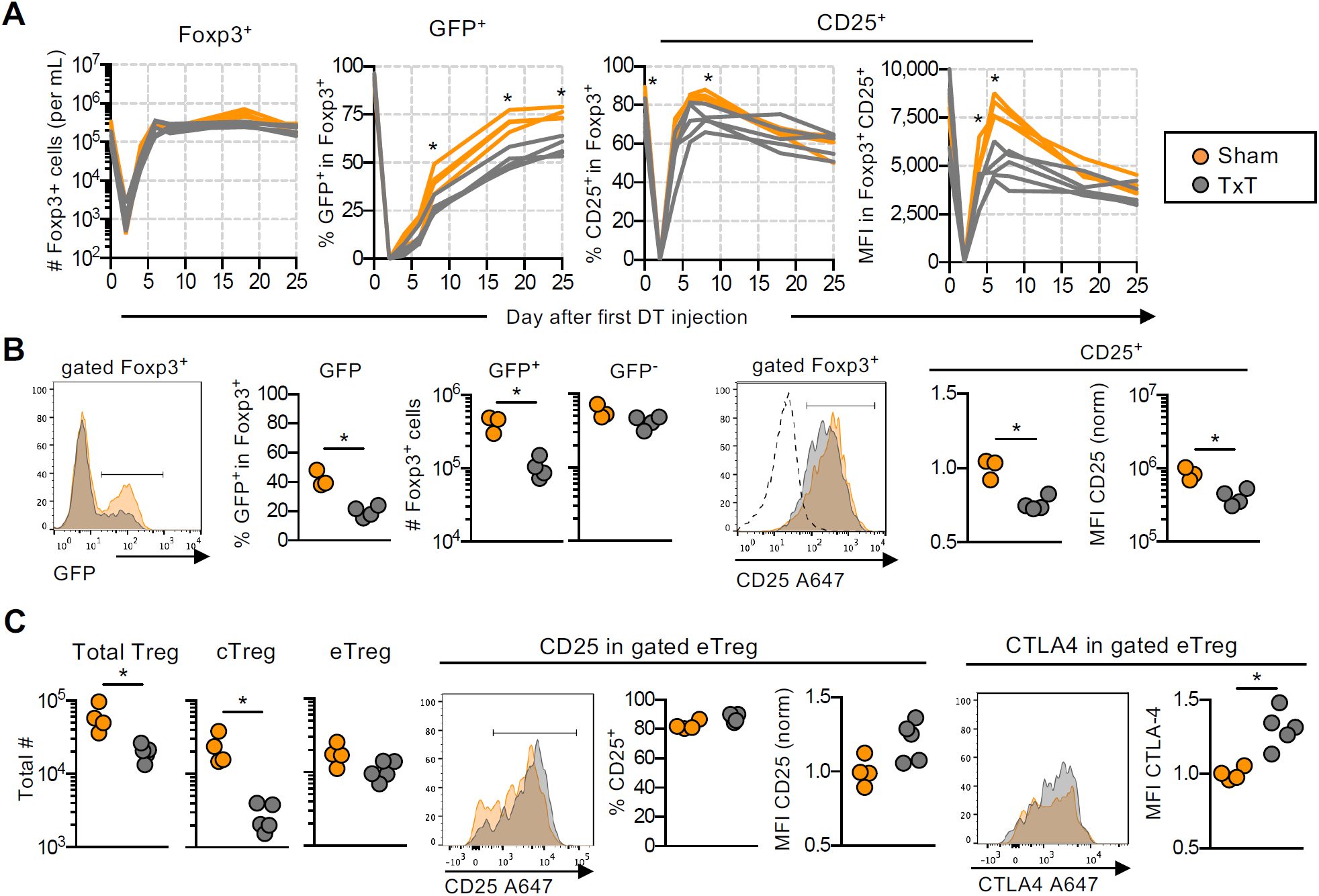
Altered return to homeostasis upon Treg depletion in thymectomized mice. Sham and TxT DEREG mice received diphtheria toxin (DT) on day 0 and 1. **A**) Peripheral blood samples were collected and analyzed on day 0, 2, 4, 6, 8, 18 and 25. Shown left to right is the analysis of Foxp3^+^ Treg for total numbers, frequency of GFP^+^ BAC expressors, frequency of CD25^+^ expressors, and the level of CD25 expression (MFI). Representative of two independent experiments (n=3-5). **B**) Analysis of inguinal LN on day 6 post-transfer showing representative flow cytometry plots, frequency and absolute numbers of GFP^+^ and GFP^-^ Foxp3^+^ Treg, and number of CD25+ Foxp3 Treg and their CD25 expression level (MFI normalized to average Sham values). One experiment (n=3-4). **C**) Analysis of Foxp3^+^ Treg in inguinal LN on day 35 post DT administration. Left to right, total number of Treg, CD62^+^CD44^-^ cTreg, CD62^-^CD44^+^ eTreg; representative flow cytometry plots, frequency of CD25^+^ cells and normalized MFI in eTreg; representative flow cytometry plots and normalized CTLA4 MFI in eTreg. One experiment (n=4-5). Statistical analysis was performed using the Holm-Sidak multiple comparison test for kinetics (A) and nonparametric Mann Whitney T-test for LN (B, C). *P < 0.05.

### Adult thymectomy does not prevent immune surveillance of tumor

To directly test the functional outcome of adult thymectomy on tumor growth we first monitored the CT26 cell line. This colon carcinoma is not spontaneously immunogenic but sensitive to both Treg depletion and anti-CTLA4 therapies (21, 22), making it an ideal model to test the impact of subtle immune alterations. CT26 tumor cells were implanted subcutaneously (s.c.) in the flank of syngeneic (Balb/c) alymphoid (RAG^-/-^), or wild type (WT) and DEREG mice, either TxT or Sham. Due to the heterogeneous shapes of each tumor growth curve inside each experimental group, we relied on the modified individual tumor control index (iTCI) (23, 24) which informs on tumor regression, stability and rejection, to infer rigorous statistical analysis. We first confirmed that CT26 tumors grow equally well in RAG2^-/-^ (RAG) or WT animals, and further evidenced that TxT performed 7-8 weeks before tumor implantation neither facilitated nor impaired tumor growth (**Fig. 4A**). Depletion of Treg in DEREG mice 12-13 days post CT26 implantation induces either full rejection in all animals, or prevents tumor growth in only a fraction of mice, according to the dose (1µg twice vs 0,5µg once) of administrated DT (24, 25). Testing the most sensitive protocols we evidenced that at least 80% of the mice fully rejected the tumor whether in the TxT or the Sham groups (**Fig. 4B**). Finally, when treating WT mice with anti-CTLA4 mAb during the first week after CT26 implantation, the majority of animals showed reduced or fully controlled tumor growth, irrespective of whether they were TxT or Sham (**Fig. 4C**). Together, these data establish that anti-tumor immunity is not compromised in thymectomized animals.

**Figure 4.**
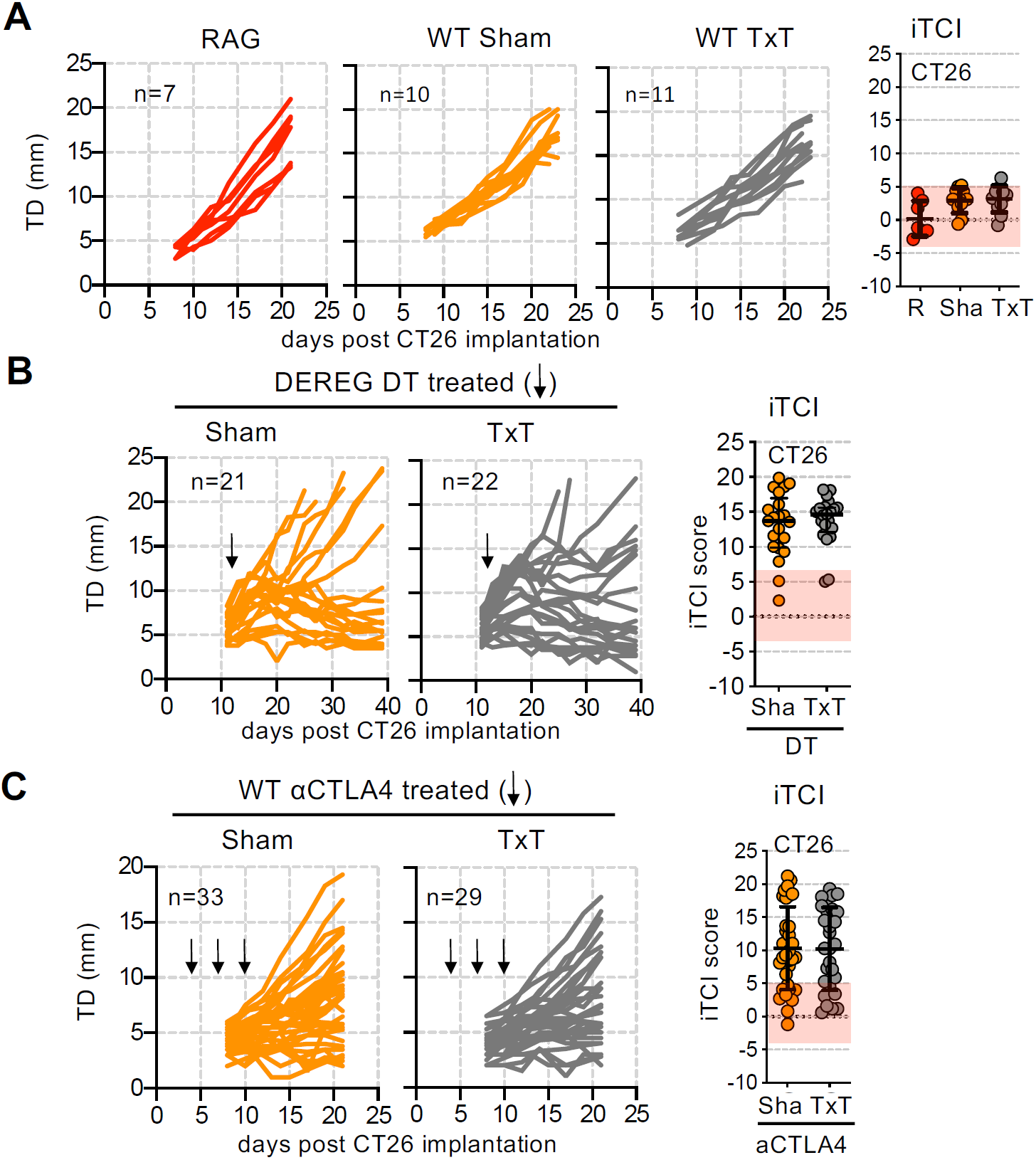
Adult thymectomy does not prevent immune surveillance of tumor. 3×10^5^ CT26 were injected s.c. in the right flank of BALB/c mice and tumor diameter (TD) in individual mice was monitored every 2-3 days. Surgery in TxT and Sham animal took place 7-8 weeks earlier. Individual Tumor Control Index scores (iTCI) performed as described in material and methods, are shown in the right panel for each experiment, the red shaded area represents the range in the RAG group. **A**) Tumor growth in alymphoid Rag^-/-^ (RAG) and TxT or Sham lymphoreplete WT mice. RAG, one experiment (n=7); TxT and Sham pool of 2 experiments (n=4-6 each). **B**) Tumor growth in diphtheria toxin (DT) treated TxT or Sham DEREG mice. The black arrow indicates the time of of DT i.p. injection (0.5ug on day 12 post tumor implantation). Pool of 4 experiments (n=4-8 each). **C**) Tumor growth in anti-CTLA4 (αCTLA4) treated TxT and Sham WT mice. Black arrows indicate time of intraperitoneal αCTLA4 administration (100 µg on days 4,7 and 10 post tumor implantation). Pool of 3 experiments (n=8-15 each). Statistical analysis for iTCI score was performed using Kruskall-Wallis multiple comparison test followed by a nonparametric Mann-Whitney test.

### Interruption of thymic activity potentiates immunotherapy of tumors expressing either weak or strong neoantigens

We next tested whether tumor intrinsic immunogenicity conditions the impact of TxT on immune surveillance. The B16F10 melanoma (here after B16) expresses several neo-antigens but remains poorly immunogenic (26) unless Treg are strongly depleted from tumor implantation on (27). The engineered B16-cOVA expresses in addition the poorly immunogenic GFP protein and the cytoplasmic portion of ovalbumin which serves as a proxy of a strong immunogenic tumor neo-antigen **(Fig. S3)**. The growth of B16-cOVA but not of B16 was equally impaired or delayed in syngeneic (C57Bl/6) TxT or Sham mice, as compared to alymphoid RAG animals (**Fig. 5A)**. We tested DT effect 13 days post-B16 tumor cells implantation into DEREG mice. In this setting, most Sham mice (76%) presented tumor growth curves similar to those of WT control animals, despite DT treatment. In contrast, a sizeable fraction of TxT hosts (70%) exerted a robust control of tumor growth upon DT treatment, resulting in higher survival (70 versus 30%) and higher iTCI scores, when compared to Sham animals (**Fig. 5B**).

**Figure 5.**
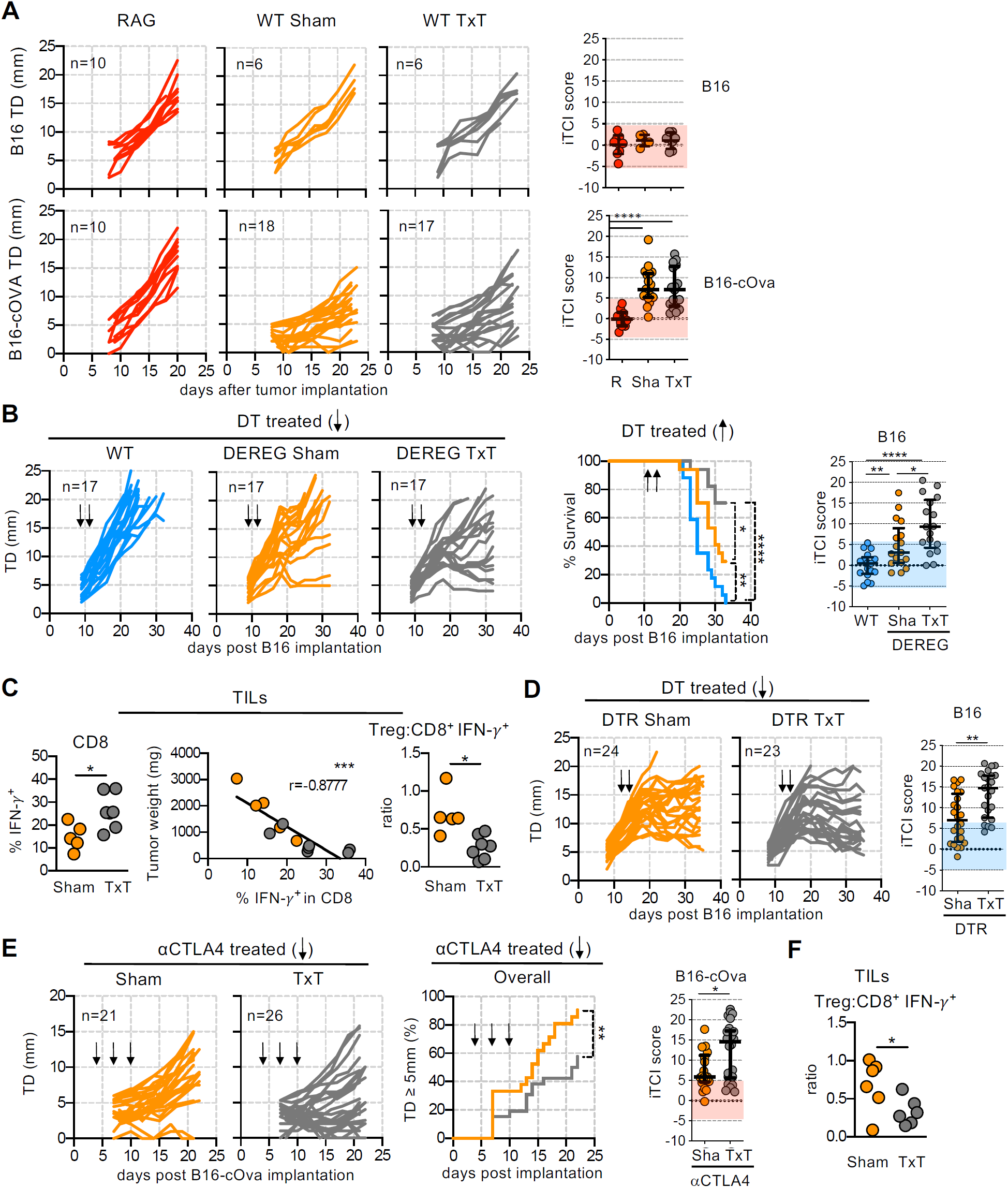
Adult thymectomy potentiates induced rejection of neoantigen expressing tumors. 2×10^5^ B16 or B16-cOVA cells were injected s.c. in the right flank of C57Bl/6 (B6) animals and tumor growth was monitored as in Fig. 4. Surgery in TxT and Sham animal took place 7-8 weeks earlier. The shaded area in the iTCI panels represent the value range in RAG (red) or WT (blue) mice. **A**) RAG and TxT or Sham lymphoreplete WT mice. Pool of 2 experiments (RAG B16 and B16-cOva, n=4-6 each; Sham and TxT B16-cOva n=2-7 each), or one experiment (Sham and TxT B16, n=6). **B**) WT and TxT or Sham DEREG, treated with DT (1µg) on days 13 and 14 post B16 tumor implantation. The central panel show mouse survival. For ethical purposes, death was defined as natural death or TD above 20mm. Pool of 2 (WT, n=3-14 each) or 3 (DEREG, n=3-8 each) experiments. **C**) B16 tumor infiltrating lymphocytes (TILs) from DT treated TxT and Sham DEREG mice analyzed 35 days post-tumor implantation. Shown are the percentage of IFN-γ producing CD8 cells (left), correlation between tumor weight and number of infiltrating IFN-γ producing CD8 cells (center) and the ratio between Treg and IFN-γ producing CD8 cells (right). One experiment (n= 5-7). **D**) DTR mice treated with DT (1µg) on days 13 and 14 post-B16 tumor implantation. Pool of 4 experiments (n=4-10). **E**) TxT and Sham WT mice bearing B16-cOVA tumors treated with αCTLA4. The central panel displays the incidence of tumor escapees given by the percentages of mice for which the TD exceeded 5mm. Pool of 4 experiments (n=3-9). **F**) B16-cOVA TILs from αCTLA4 treated TxT and Sham WT mice analyzed 12 days post-tumor implantation. Shown is the ratio between Treg and IFN-γ producing CD8 cells. One experiment (n=6). Statistical analysis for TCI score performed as in Fig 4. Statistical analysis for survival curves used log-rank test. Statistical analysis in (C and F) was performed using nonparametric Mann Whitney T-test, and Pearson correlation coefficients. *P < 0.05, **P < 0.005, and ***P < 0.001, ****P < 0.0001.

Cellular analysis of tumor infiltrating lymphocytes (TIL) by the end of the experiment (day 35) revealed a 2 fold higher frequency of CD8 cells producing IFN-γ, a parameter inversely correlated with tumor weight, and lower ratio of Treg to CD8 IFN-γ producers, a hallmark of productive antitumor immunity, in TxT when compared to Sham animals (**Fig. 5C**). Other cellular features included increased number of activated CD4 or CD8 cells and increased frequency of highly cycling and KLRG1^+^ terminally differentiated Treg in TxT vs Sham mice (**Fig. S4**). Similar enhanced control of tumor growth was revealed when testing mice carrying the DTR gene inserted at the Foxp3 locus (DTR mice). In DTR, contrarily to DEREG animals, all Treg are equally targeted upon DT administration. In this model, the vast majority of mice (96%) responded to DT treatment by arrest or regression of tumor growth, and TxT animals performed better for the latter, resulting in higher iTCI scores (**Fig. 5D**).

We next confirmed that anti-CTLA4 therapy fails to alter B16 tumor growth (22) and evidenced this feature is not altered in TxT mice (**Fig. S5**). Surprisingly, the B16-cOVA which we showed above is spontaneously immunogenic in Sham mice did not respond to the therapy. In contrast, 60% of the animals in the TxT group, compared to 20% in the Sham cohort, controlled tumor growth for at least 20 days, maintaining a tumor size lower than the threshold of precision for measurement. Analysis of tumor growth curves evidenced a striking bimodal iTCI score distribution in TxT mice (**Fig. 5E**), not related to specific experimental groups. Cellular analysis of B16-cOVA TILs confirmed a reduced ratio of Treg to CD8 IFN-γ producers in TxT when compared to Sham treated animals **(Fig. 5F**), in this case seemingly the result of decreased frequency of Treg (**Fig. S6**). We conclude that continuous activity of the thymus is not required for a positive outcome of anti-CTLA therapy and may actually be unfavorable. Together, these data confirm that adult thymectomy does not prevent tumor immune surveillance, even if induced and irrespectively of the tumor intrinsic immunogenicity, and reveals it may improve Treg based therapy.

## Discussion

In this work we document that cessation of thymic activity in adult alters Treg composition and function in the periphery. We demonstrate, across tumor models and mouse strains, that adult thymectomy *per se* does not provide for weaker antitumor immunity in mice and may actually favor the outcome of immunotherapies. To our knowledge, our work is the first to address experimentally the contribution of the thymus on tumor growth in otherwise healthy adults. Together, our findings point to a scenario in which abrogation of thymic activities affects preferentially the regulatory over the ridding arm of the immune activities elicited by tumors.

To address the impact of thymic activity decline on immune regulation, we chose adult thymectomy as a study model rather than aged animals. This strategy allows to dissociate aging of the peripheral T cell pool *per se* from other indirect effects driven by systemic aging. For instance, other subset of immune cells, including dendritic cells that are mandatory partners of T cells, do lose competences with age (28), and senescence-associated secretory phenotype (SASP) factors produced by multiple aged-tissues have systemic and pleiotropic effects. Full ablation of the thymus after production of a normal initial T cell pool is nowadays rarely performed and limited to sporadic cases of adults with myasthenia gravis and to newborn/children presenting congenital heart defects (partial or total ablation of the thymus eases the surgical access to the heart). Follow-up of these children reveals reduced naïve T cells number and TCR diversity, but not impaired immune responses to common infections and most vaccines (29). While these cohorts are small, and the follow-up time is in the range of one or two decades, it is noticeable that increased tumor incidence has not been reported.

Thymic output in mice has been studied in details, notably through the analysis of animals bearing the GFP gene inserted at the RAG locus, where GFP expression in the periphery reports T cells that recently exited the thymus. These studies established that full maturation of RTE is achieved in about 3 weeks (30) which certifies absence of RTE in our experimental animals 6-8 weeks post TxT. Kinetic analysis also established that decline in RTE number starts soon after birth and reaches its maximum by 30 weeks of age (31). At the age we analyzed mice (12-17 weeks) RTE represents about 20% of T cells corresponding to 3- 4×10^6^ cells/spleen (31), ensuring our Sham controls had an active thymus. These analyses were performed in B6 animals, and it is possible that BALB/c mice have different kinetics of RTE decline. It is also possible that the cervical thymi reported in BALB/c (32), while atrophic at steady state, compensated for, and mitigated the effect of, thoracic thymus ablation.

Our detailed cellular analysis confirms our prediction that adult thymectomy alters both Tconv and Treg composition, to somewhat similar extent, and imposes a similar bias for activated/effector/memory phenotype in the peripheral T cell pool. Our results also confirms that absence of tTreg output is not compensated by differentiation of pTreg in the periphery, as was predicted by the earlier finding that RTE are the precursors of *bona fide* Treg differentiated in the periphery (7, 8). This finding is also consistent with the compromised return to homeostasis upon Treg perturbation, which we document in TxT animals. Together, these evidences support our original proposition that predicting the impact of thymic cessation on the outcome of an immune response, and even more so for a composite of self- and nonself-antigens as for a tumor, is non-trivial.

By testing tumor cell lines of distinct origins, presenting different degree of immunogenicity, and two strains of mice, we evidence that cessation of thymic activity for 2 months does not reduce spontaneous immune surveillance or impairs immuno-therapeutic interventions. While our experimental design does not model years of aging in humans, it may model thymic atrophy induced by chemotherapy and other biological stress. We further document that in some instances TxT favors a positive outcome of immuno-therapies, likely the result of a subtle tilting in the Treg to effector balance, and not predicted by the intrinsic immunogenicity of each tumor model. As a proxy of immune therapies of cancer in humans, we used induced depletion of Treg and anti-CTLA4 treatment. The immune checkpoint inhibitor anti-CTLA4, was demonstrated to kill Treg in vivo (33) and the higher the efficacy in Treg killing the better the therapeutic outcomes (34). Despite clear improvement in survival rate, some patients do not respond to such therapies and others fail to achieve a complete response; a variation not yet understood. A striking feature of our data is actually the heterogeneity of tumor growth kinetics in a single experimental group, observed only in situation of spontaneous or induced immunogenicity, and despite the experimental paradigm of tumor cell lines and isogenic mice to reduce biological variation. It would appear that interindividual variation in such restricted setting remains to be formally addressed before one could approach the full complexity of human heterogeneity.

Studies and reviews addressing the impact of thymic involution or ablation on health still reduce T cells to a pool of effector cells and appear oblivious to more than two decades of evidences that Foxp3^+^ Treg are essential to ensure “dominant” tolerance. In contrast, Treg-based therapies are in the clinic or being developed to the benefit of autoimmunity and cancer (9, 35). In agreement with the notion that thymic senescence affects Treg are the epidemiological evidence that while most autoimmune diseases first manifest in children and young adults, a second wave of incidence is noticeable in older individual, when thymic activities have significantly declined. Moreover, TxT NOD mice developed diabetes strikingly faster than Sham controls (36), and corticosteroid treatment which induces thymocytes death further enhances the effects of cyclophosphamide-induced Treg depletion (37).

Together our analysis points to a scenario by which reduction in thymic output, be it the result of surgical ablation, stress induced contraction or age-related involution has a dual effect altering not only effector but also Treg (see **Figure 6**). As each cell compartment encompasses a very large repertoire of different TCR sequences, although larger for effector cells than for Treg, the reservoir of reactivities in each compartment is mildly affected by interrupted thymic output. In situation of sustained cessation of thymic output, the reduction in the TCR repertoire of both effector and Treg cells affects the domain of competences of each cell subset (ridding and regulation, respectively) which in most cases results in unperturbed interactions between these two compartments, and thus preserves tolerance to self-tissues and tumors. Upon added stress on the Treg compartment, such as during immune check-point inhibitor therapy, the inability of the thymus to replenish this cell pool results in unbalanced immune responses, including enhanced anti-tumor immunity. This model highlights the robustness of the immune system, explains the overall efficient immune responses (29) and mild increase in auto-reactive Ab (38) presented by humans submitted to surgical thymectomy, and argues that higher incidence of tumors with age cannot be solely attributed to thymic output decline. Further refinement of this scenario awaits technological development allowing extensive and unbiased reactive TCR repertoire analysis.

**Figure 6.**
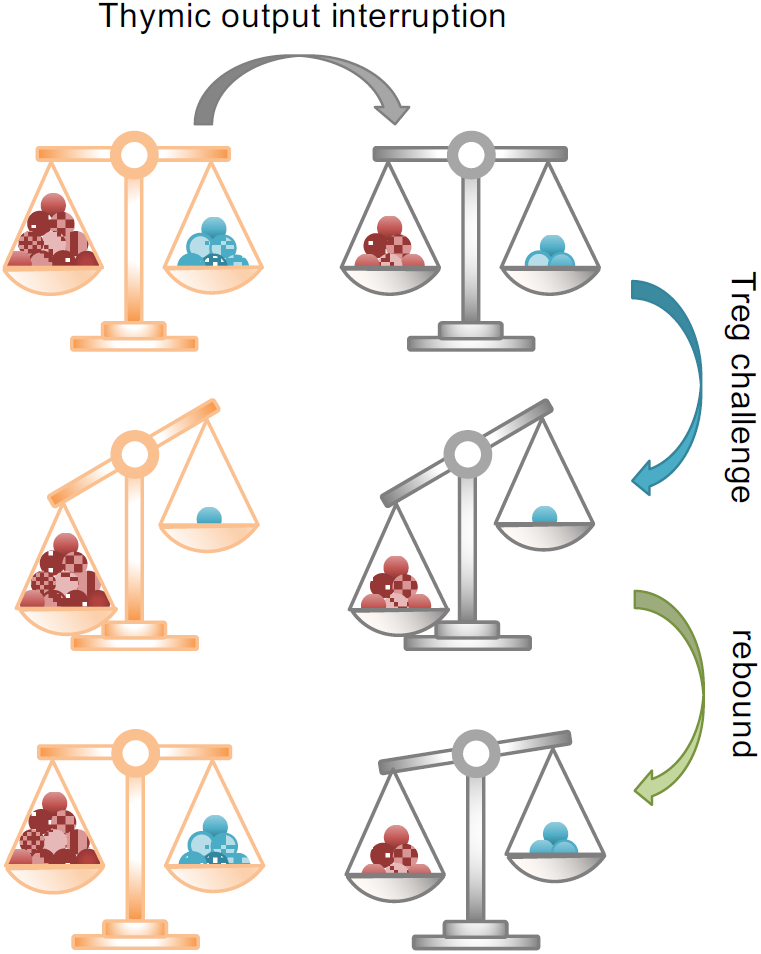
schematic representation of the model. Effector T cells and Treg are represented by red and blue filled circles, respectively. Equilibrium at steady state (upper left) is the result of cell numbers and cell composition at the reactive repertoire level. Adult thymectomy alters equally Tconv and Treg numbers and composition (upper right). Diphtheria toxin in DEREG or DTR mice or anti-CTLA4 mAb in WT animals decreases Treg number unleashing effector cells (middle panels). Thymic output restores Treg numbers and diversity (lower left). In absence of thymic activity, the rebounding Treg have limited diversity and unmatched repertoire results in subtle disbalance (lower right).

## Materials and Methods

### Mice

C57BL/6J, C57BL/6.Thy1.1 congenic (Jackson), C57BL/6-Tg(Foxp3-DTR/EGFP)23.2Spar/Mmjax (B6 DEREG), B6.129(Cg)-Foxp3tm3(DTR/GFP)Ayr/J (B6 DTR), C57BL/6.RAG2.KO, C57BL/6.TCRβ.KO, C57BL/6.Foxp3^tm2Ayr^ (Foxp3^fGFP^) C57BL/6.Foxp3^tm1(CD2/CD52)Shori^.CD45.1.DEREG (B6. Foxp3^**hCD2**^.CD45.1 DEREG), BALB/c.ByJ, BALB/c-Tg(Foxp3-DTR/EGFP) Spar/Mmjax (Ba DEREG), and BALB/c.RAG2.KO mice were bred and raised at the Instituto Gulbenkian de Ciência (IGC) animal facility under specific pathogen– free conditions. Thymectomies (TxT) were performed in aseptic conditions on 5-7 weeks old mice under ketamine/xylazine anesthesia. Sham-thymectomy (Sham) was complete surgical procedure without removal of thymic lobes. Procedures were conducted according to the Federation for Laboratory Animal Science Association guidelines and approved by the ethical committee of the IGC and the Portuguese state institution (DGAV).

### Cell lines and tumor models

Tumor cells were cultured at 37°C in RPMI 1640 (Life Technologies), supplemented with 10% Fetal Bovine Serum (South America), Premium (Biowest), 1% Penicillin-Streptomycin (Life Technologies), 0.1% Gentamicin (Life Technologies) and 0.1% 2-Mercaptoethanol (Life Technologies). To prepare for tumor cell injections, cells were trypsinized (Life Technologies) and ressuspended in ice cold HBSS with no calcium and no magnesium (Life Technologies). Mice were injected subcutaneously, in the right flank, with 2×10^5^ B16-F10-luc2 (B16) (CaliperLS), or with B16-cOVA. The latter were B16-F10-luc2 transduced with a GFP-ovalbumin vector (Nir Hacohen) (**Fig. S3**) kindly offered by Catarina Moita. The B16-cOVA cell line were sorted by flow cytometry to select GFP^+^ cells, and cultures regularly checked to guaranty >98% GFP^+^ cells. For CT26 (ATCC) experiments, mice were injected with 3×10^5^ tumor cells per mouse. Tumor size was monitored with a caliper every 2-3 days from day 8 days post tumor injection. Tumor Diameter (TD) was calculated as TD= (L+W)/2. For ethical reasons mice were sacrificed when TD≥20mm. Where indicated, mice received an i.p. injection with DT (0,5 or 1µg per injection) (322326-1, Calbiochem) diluted in 100 µl PBS on day 13 and 14 after tumor injection, or with 100 ul of PBS containing 100 ug of anti-CTLA4 monoclonal antibody (clone 4F10, produced in house) on day 4 and 7 and 10 after tumor cells injection.

### Flow cytometry analysis

Single-Cell suspensions were prepared from harvested mouse inguinal lymph nodes (iLN) by mechanical disruption using a buffer containing 10% PBS (Life Technologies) and 2% Fetal Bovine Serum (South America), Premium (Biowest). As for tumor infiltrating lymphocytes (TIL) isolation, tumors were digested in HBSS medium (Life Technologies) containing EDTA (Gibco), BSA (Sigma), collagenase type IV (Sigma) and DNAse I (Sigma) for 30 minutes after which a Percol gradient (Sigma) was used to isolate lymphocytes. Live cells were counted using magnetic beads and Propidium Iodide. To stimulate the production of cytokines, cells were incubated for 4h at 37°C in RPMI 1640 (Life Technologies) containing PMA (phorbol 12-myristate-13-acetate) (Sigma), Ionomycin (Sigma) and protein transport inhibitor (Brefeldin A) (eBioscience). After pre-incubation with Fc-block, surface markers were stained using antibodies described in table S1. Cells were then incubated overnight at 4°C in Foxp3 fix/permeabilization buffer (eBioscience) and stained for intracellular transcription factors and cytokines using antibodies also described in table S1. For analysis of GFP expressing cells in DEREG animals, the GFP signal was enhanced by using anti-GFP antibody staining. Analysis was performed using Cyan ADP or BD Fortessa X20. Data were analyzed with Flowjo.

### Adoptive cell transfer

Specific cell subsets were isolated by flow cytometry and transferred by retroorbital injection. When indicated, recipient mice were treated with DT (1µg in 100 µl) one day before an at the day of cell transfer. To assess Treg stability and suppressive function in vivo, CD4^+^CD8^-^GFP^+^ (Treg), CD4^+^CD8^-^GFP^-^ (CD4), and CD4^-^CD8^+^ (CD8) cells isolated from pooled spleenocytes and peripheral lymph node cells of congenic Thy 1.1 and Thy 1.2 Foxp3^fGFP^ reporter mice. 1×10^5^ Treg cells and 4×10^5^ CD4 or CD8 cells were transferred into TCRβ ^−/−^ recipients. For in vivo pTreg differentiation, 3×10^5^ CD4^+^CD8^-^GFP^-^CD44^-^ naïve cells isolated from peripheral lymph node cells of Thy 1.1 Foxp3^fGFP^ reporter mice were transferred into congenic Thy 1.2 wild type and DT treated DEREG mice. To assess GFP expression in DEREG pTreg, 3×10^5^ CD4^+^CD8^-^ hCD2^-^CD44^-^ naïve thymocytes isolated from CD45.1 DEREG Foxp3^hCD2^ reporter mice were transferred into congenic DT treated CD45.2 DEREG recipients. To test for GFP acquisition by DEREG GFP^-^ Treg, CD4^+^CD8^-^hCD2^+^GFP^-^ cells were isolated from pooled splenocytes and peripheral lymph node cells of CD45.1 DEREG Foxp3hCD2 mice that were DT treated 5 days before, and transferred into DT treated congenic CD45.2 DEREG. Cell sorting was performed on BD FACS Aria.

## Statistical analysis

The individual Tumor Control Index (iTCI) was derived from the TCI (23) which compiles three scores per experimental group assessing tumor inhibition, stability and rejection. We modified this method to calculate the three sub-scores for each individual mouse (iTCI), an improvement which allows for statistical analysis between experimental groups (24). Statistics of iTCI was performed using Kruskall-Wallis multiple comparison test followed by a nonparametric Mann-Whitney test. Statistics of all cellular analysis were performed using nonparametric Mann-Whitney test. Logrank tests were used for survival curves. Correlation analyses were performed using Pearson correlation coefficients.

## Acknowledgements

We are grateful to the core facilities at IGC: Ana Regalado for mAb production and the teams Marta Monteiro (flow cytometry) and Manuel Rebelo (mouse husbandry). We thank Karine Serre, Ricardo Paiva and Iris Caramalho for critical discussions during the conduction of this work, and Luis Moita and Catarina Moita for providing the B16-cOVA line. This work was supported by the IGC-Fundação Calouste Gulbenkian and by the Portuguese scientific council (Fundação para a Ciência e a Tecnologia) including fellowships to JGS (SFRH/BD/52435/2013), IC (SFRH/BPD/111454/2015) and MLB (283/BI/15; UID/Multi/04555/2013). Mouse experiments were also in part supported by the national infrastructure CONGENTO LISBOA-01-0145-FEDER-022170 (FCT, Lisboa2020, Por2020, ERDF). JAS, MLB and JD designed the research, JAS, MLB and IAC performed experiments, JAS, MLB and JD analyzed the data, JAS and JD wrote the manuscript.

## Conflicts of interest declaration

The authors declare no commercial or financial conflict of interest.

## Supplementary material

**Supplementary Table S1.** monoclonal Antibodies used in this study.

| Reactivity | Clone | Conjugate | Supplier |
| --- | --- | --- | --- |
| CD16/CD32 (Fc-block) | 2.4G2 | *none* | In house |
| CD4 | GK1.5 | PerCP-Cy5.5 / PE-Cy7 | Biolegend |
| CD8 | YTS169.4 | FITC/Biot/PB/A647 | In house |
| CD25 | PC-61 | Biot/A647 | In house |
| CD44 | IM7 | FITC/Biot. | BD Pharmingen |
|  |  | e450 | eBioscience |
| CD45.1 | A20 | PE | Biolegend |
|  |  | PB | In house |
| CD45.2 | 104.2 | A647 | In house |
| CD62L | MEL-14 | FITC/Biot/PE | eBioscience |
| CD152 (CTLA-4)<br>* | 4F10 | *none* | In house** |
|  |  | A647 | In house |
|  | UC10-4B9 | PE | eBioscience |
| Foxp3* | FJK-16s | FITC/e450/APC | eBioscience |
| GFP | 19C8 | A488 | In house |
| Helios* | 22F6 | PE | Biolegend |
| Human/NHP CD2 | RPA-2.10 | APC | eBioscience |
| IFN-γ* | XMG1.2 | PE | BD Pharmingen |
| Ki67* | SolA15 | FITC/A700 | eBioscience |
| KLRG1 | 2F1 | APC | BioLegend |
| Neuropilin-1 | Polyclonal | APC | R&D |
| Qa-2 | 1-1-2 | Biot | BD Pharmingen |
| Streptavidin | n.a. | APC-Cy7 | BioLegend |
| TCRβ | H57-597 | PE | BioLegend |
|  |  | PerCP-Cy5.5 | eBioscience |
|  |  | A647 | In house |
| TIGIT | 1G9 | PE | BioLegend |
| Thy 1.1 | OX-7 | Biot. | BioLegend |
| Thy 1.2 | 30H12 | APC | In house |
| TNF-α* | MP6-XT22 | PE | BioLegend |

A, Alexa fluor; APC, allophycocyanin; Biot, biotin; Cy, cychrome; PB, pacific blue; PE, phycoerythrin; FITC, fluorescein isothiocyanate; e, eFluor; PerCP, peridinin-chlorophyll-protein
\* intracellular/intranuclear staining
\*\*All antibodies produced in house were purified on protein G columns. When indicated Ab were also conjugated in house

**Figure S1.**
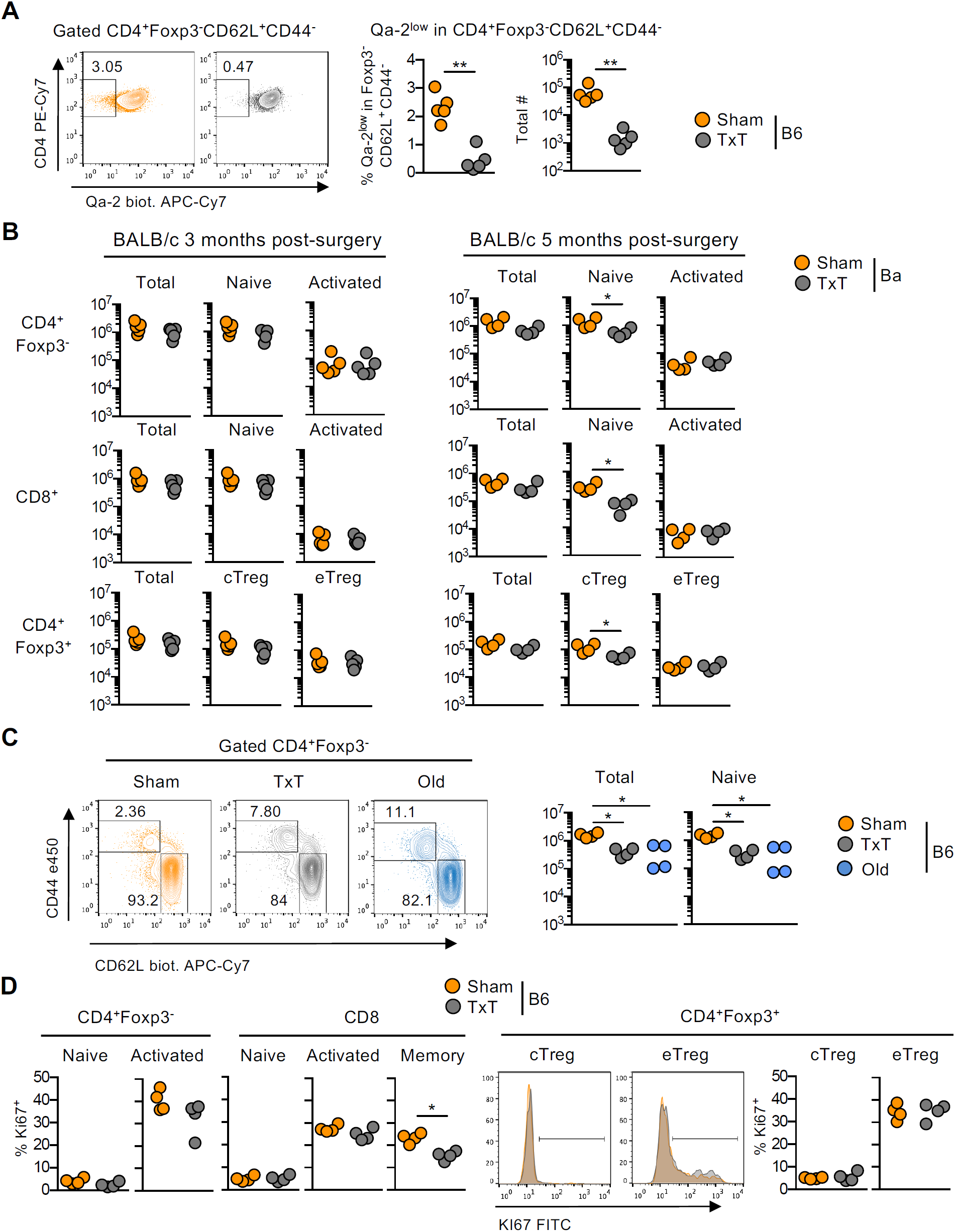
Complement to Figure 1. Lymph nodes from B6 (A, C, D) or BALB/c (B) TxT and Sham mice prepared as in Fig. 1, or from 15-month-old unmanipulated B6 animals were analyzed by flow cytometry. **A**) Representative flow cytometry analysis and compiled data for frequency and number of recent thymic emigrants (RTE) identified as CD4^+^Qa-2^low^Foxp3^-^CD62L^+^CD44^-^ in B6 mice **B**) Analysis of BALB/c mice. Surgery was performed at 5/6 weeks of age and analysis conducted 3 (**left**) or 5 (**right)** months later. Shown are number of total, naïve and activated CD4 and CD8 T cells, as well as Treg, cTreg and eTreg, defined by CD44 and CD62L expression as in Fig. 1. **C**) Activated and naïve CD4 T cells as defined in Fig.1, comparing Sham, TxT and old B6 animals. **D**) Frequency of Ki67 expressing cells in Foxp3^-^CD4^+^ T cells (left), CD8 T cells defined here as TCRβ^+^ CD4^-^ (right), and Foxp3^+^ CD4^+^ Treg (bottom). Each cell subset was further subdivided using the CD44 and CD62L markers, as in Fig. 1. One experiment (n=4-5). Statistical analysis was performed using nonparametric Mann Whitney T-test. *P < 0.05.

**Figure S2.**
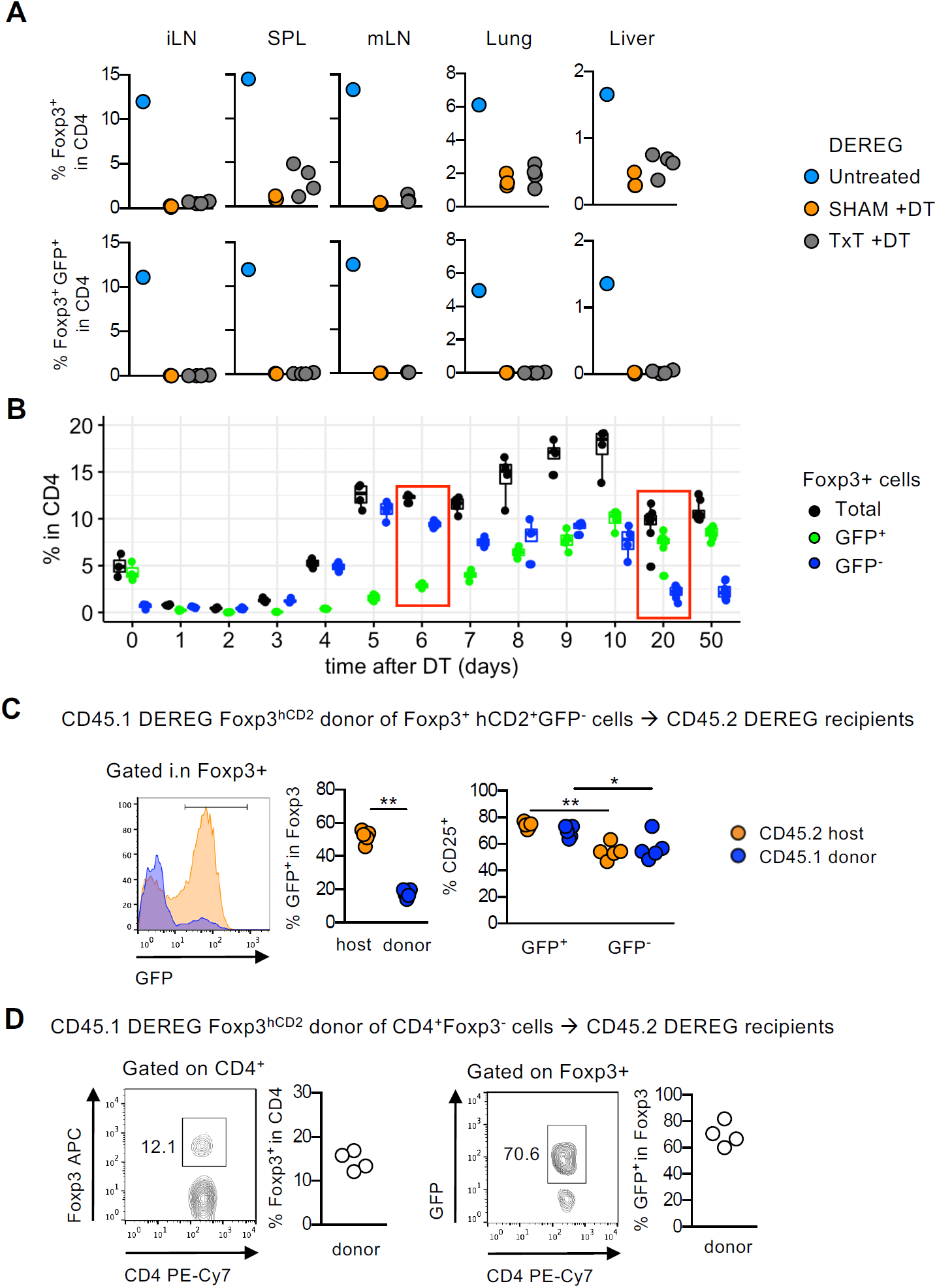
Unlike newly differentiated pTreg or tTreg, GFP^-^ Treg in DEREG mice are not precursors of *bona fide* GFP^+^ Treg. **A**) Efficiency of Treg depletion in DEREG mice. Cellular analysis of untreated euthymic, TxT and Sham B6 DEREG mice 3 days after the first diphtheria toxin (DT) administration (1ug per injection on two consecutive days). Shown are the percentages of CD4 cells that are Foxp3^+,^ and the frequency of Foxp3^+^ cells that express the BAC reporter (GFP^+^). LN: inguinal lymph node, SPL: spleen, mLN: mesenteric lymph node. One experiment (n=3-4). **B**) Kinetic of Treg rebound in B6 DEREG mice. Frequency of total, BAC-GFP^+^ and BAC-GFP^-^ Foxp3^+^ CD4 cells in peripheral blood samples from DT treated DEREG mice. Two groups of mice (n=4 each) were followed on alternate days post-DT injection. One representative of three independent experiments. Red frames indicate the days selected in Fig. 3 B, C for lymphoid organs analysis. **C**) GFP conversion experiment. hCD2^+^ GFP^-^ CD4 cells were isolated from pooled spleen and lymph node of B6 CD45.1 huCD2 DEREG mice and transferred into B6 CD45.2 DEREG recipients. Donors were treated with DT 5 days earlier, recipients 1 day earlier and at the day of transfer. Analysis of donor and recipient Foxp3^+^ cells was performed 1 week post-transfer. Shown are a representative flow cytometry plot, used to infer the frequency of BAC expressors GFP^+^ Treg from each origin, as well as the frequency of CD25^+^ cells in BAC-GFP^+^ and BAC-GFP^-^ Treg. One experiment (n=5). **D**) Foxp3 conversion experiment. CD45.1^+^CD4^+^CD8^-^Foxp3^-^ thymocytes isolated from B6 CD45.1 huCD2 DEREG donors were transferred into B6 CD45.2 DEREG recipients that were treated with DT 1 days earlier and at the day of transfer. Analysis of donor cells was performed 1 week post-transfer. Shown is representative flow cytometry analysis, the percentages of Foxp3^+^ (pTreg) inside CD45.1^+^ CD4 donor cells, and the frequency of BAC-GFP expressors inside newly differentiated pTreg. One experiment (n=4). Statistical analysis was performed using nonparametric Mann Whitney T-test. *P < 0.05, **P < 0.005.

**Figure S3.**
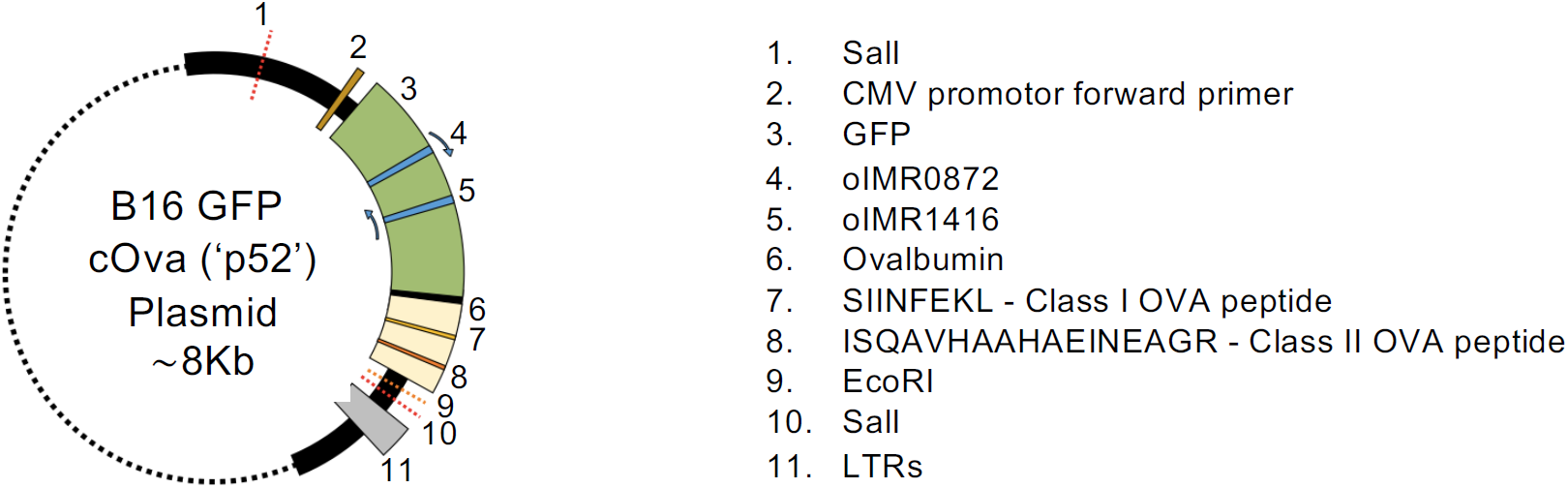
Complement to Figure 5A. Schematic representation of the c-OVA construct. Sanger sequencing confirmed the ovalbumin protein is truncated excluding the secreted and transmembrane part of the molecule, and containing only the intra cytoplasmic domain.

**Figure S4.**
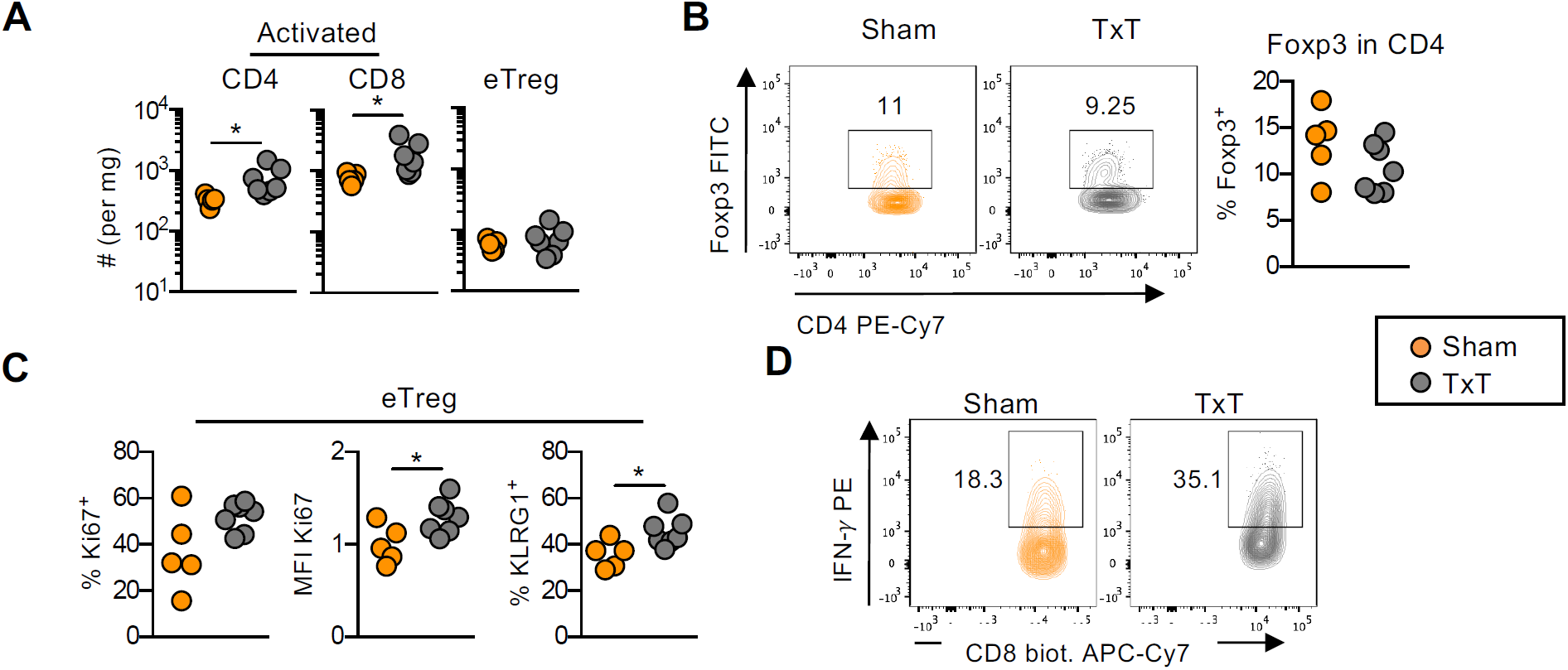
Complement to Figure 5C. B16 TILs from DT treated TxT and Sham B6 DEREG mice analyzed 35 days post-tumor implantation. **A**) Total numbers of activated (CD44^+^ CD62L^-^) Foxp3^-^ CD4^+^ cells, CD8 cells and Foxp3^+^ CD4^+^ eTreg cells per mg of tumor. **B**) Representative Flow cytometry analysis and percentage of Foxp3^+^ CD4 cells. **C**) Shown is an analysis of CD44^+^ CD62L^-^ Foxp3^+^ eTreg for frequency of Ki67^+^, MFI of Ki67 in Ki67^+^ cells, and percentage of KLRG1^+^ cells. MFI: mean fluorescence intensity (normalized to average Sham values). **D**) Representative flow cytometry analysis to identify Interferon-*γ*(IFN-γ) producing CD8 cells, percentage shown in Fig. 5C. One experiment, Statistical analysis was performed using nonparametric Mann Whitney T-test. *P < 0.05

**Figure S5.**
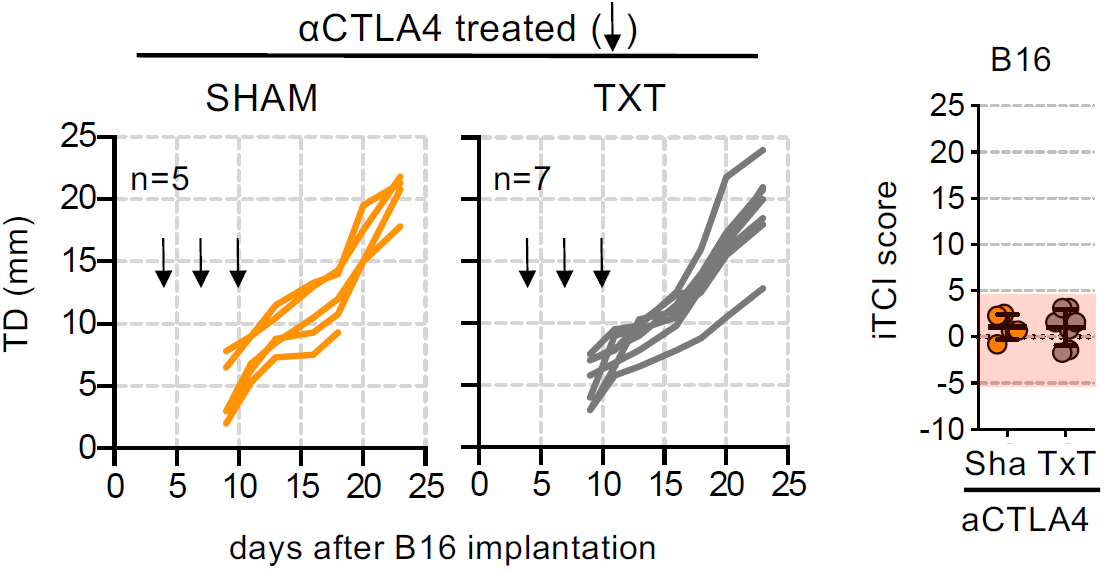
Complement to Figure 5E. 2×10^5^ B16 cells were implanted s.c. into TxT and Sham B6 mice treated with αCTLA4. Black arrows indicate time of i.p. administration of αCTLA4 (100ug on days 4, 7 and 10). One experiment (n=5-7). Statistical analysis for iTCI score performed using nonparametric Mann-Whitney test, not significant.

**Figure S6.**
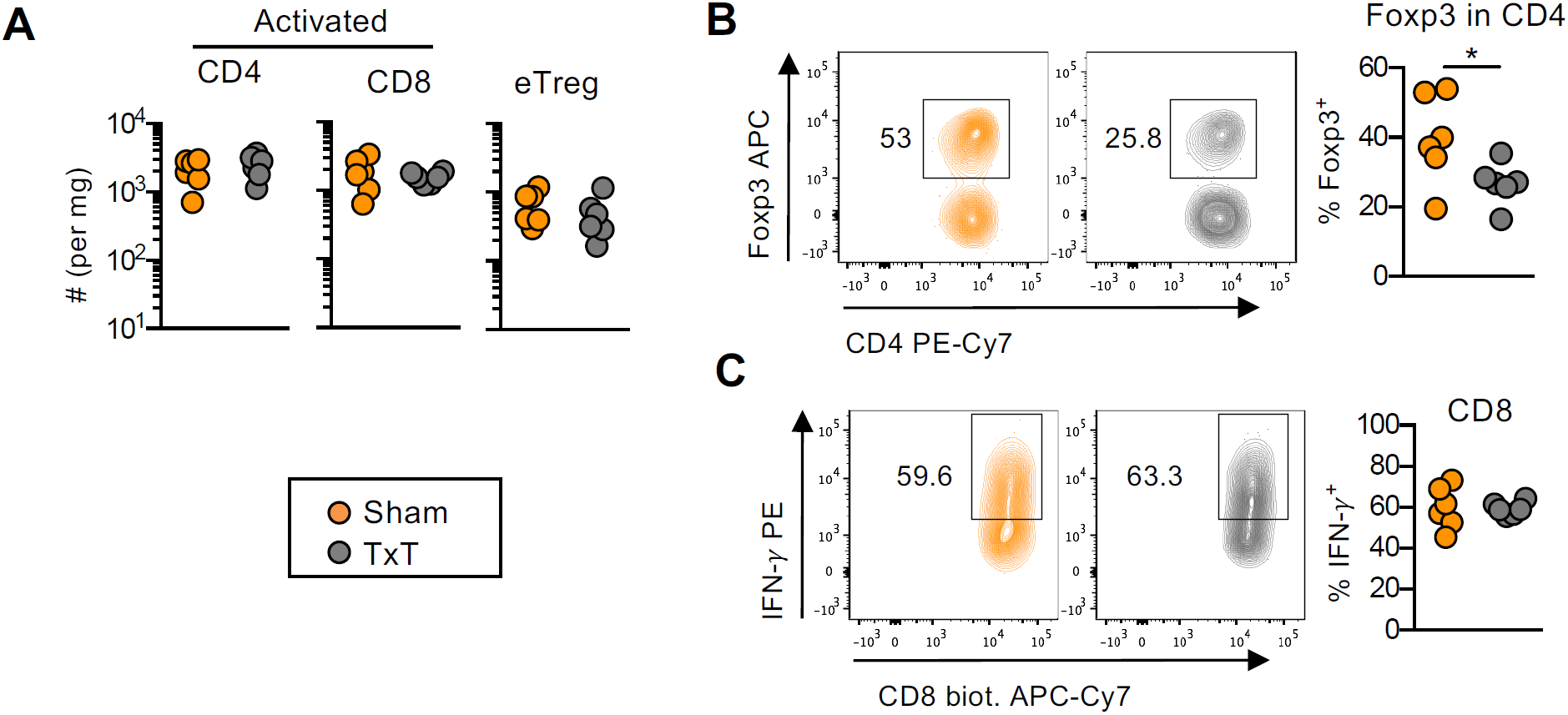
Complement to Figure 5F. B16-cGFPOVA tumor infiltrating lymphocytes from αCTLA4 treated TxT and Sham B6 mice were analyzed 12 days post-tumor implantation. **A**) Total numbers of activated (CD44^+^ CD62L^-^) Foxp3^-^ CD4^+^ cells, CD8 cells and Foxp3^+^ CD4^+^ cells per mg of tumor. **B**) Representative flow cytometry analysis and percentage of Foxp3^+^ cells inside gated CD4 T cells. **C**) Representative flow analysis and percentage of Interferon-*γ*(IFN-γ) producing CD8 T cells. One experiment (n=6). Statistical analysis was performed using nonparametric Mann Whitney T-test. *P < 0.05.

## References

1. A. Y. Rudensky, Regulatory T cells and Foxp3. Immunol Rev 241, 260–268 (2011).

2. S. Sakaguchi, K. Wing, Y. Onishi, P. Prieto-Martin, T. Yamaguchi, Regulatory T cells: how do they suppress immune responses? Int Immunol 21, 1105–1111 (2009).

3. R. Bacchetta, F. Barzaghi, M. G. Roncarolo, From IPEX syndrome to FOXP3 mutation: a lesson on immune dysregulation. Ann N Y Acad Sci 1417, 5–22 (2018).

4. C. S. Hsieh et al., Recognition of the peripheral self by naturally arising CD25+ CD4+ T cell receptors. Immunity 21, 267–277 (2004).

5. J. Carneiro et al., When three is not a crowd: a Crossregulation model of the dynamics and repertoire selection of regulatory CD4+ T cells. Immunol Rev 216, 48–68 (2007).

6. S. Zelenay et al., Cutting edge: Intrathymic differentiation of adaptive Foxp3+ regulatory T cells upon peripheral proinflammatory immunization. J Immunol 185, 3829–3833 (2010).

7. R. S. Paiva et al., Recent thymic emigrants are the preferential precursors of regulatory T cells differentiated in the periphery. Proc Natl Acad Sci U S A 110, 6494–6499 (2013).

8. A. Pratama, A. Schnell, D. Mathis, C. Benoist, Developmental and cellular age direct conversion of CD4+ T cells into RORgamma+ or Helios+ colon Treg cells. J Exp Med 217 (2020).

9. A. Tanaka, S. Sakaguchi, Regulatory T cells in cancer immunotherapy. Cell Res 27, 109–118 (2017).

10. H. Gogas et al., Prognostic significance of autoimmunity during treatment of melanoma with interferon. N Engl J Med 354, 709–718 (2006).

11. S. Das, D. B. Johnson, Immune-related adverse events and anti-tumor efficacy of immune checkpoint inhibitors. J Immunother Cancer 7, 306 (2019).

12. A. Snyder et al., Genetic basis for clinical response to CTLA-4 blockade in melanoma. N Engl J Med 371, 2189–2199 (2014).

13. S. Malchow et al., Aire-dependent thymic development of tumor-associated regulatory T cells. Science 339, 1219–1224 (2013).

14. R. Alonso et al., Induction of anergic or regulatory tumor-specific CD4(+) T cells in the tumor-draining lymph node. Nat Commun 9, 2113 (2018).

15. S. Palmer, L. Albergante, C. C. Blackburn, T. J. Newman, Thymic involution and rising disease incidence with age. Proc Natl Acad Sci U S A 115, 1883–1888 (2018).

16. K. S. Smigiel et al., CCR7 provides localized access to IL-2 and defines homeostatically distinct regulatory T cell subsets. J Exp Med 211, 121–136 (2014).

17. I. den Braber et al., Maintenance of peripheral naive T cells is sustained by thymus output in mice but not humans. Immunity 36, 288–297 (2012).

18. E. Szurek et al., Differences in Expression Level of Helios and Neuropilin-1 Do Not Distinguish Thymus-Derived from Extrathymically-Induced CD4+Foxp3+ Regulatory T Cells. PLoS One 10, e0141161 (2015).

19. J. H. Duarte, S. Zelenay, M. L. Bergman, A. C. Martins, J. Demengeot, Natural Treg cells spontaneously differentiate into pathogenic helper cells in lymphopenic conditions. Eur J Immunol 39, 948–955 (2009).

20. S. Rausch et al., Establishment of nematode infection despite increased Th2 responses and immunopathology after selective depletion of Foxp3+ cells. Eur J Immunol 39, 3066–3077 (2009).

21. S. A. Fisher et al., Transient Treg depletion enhances therapeutic anti-cancer vaccination. Immun Inflamm Dis 5, 16–28 (2017).

22. J. F. Grosso, M. N. Jure-Kunkel, CTLA-4 blockade in tumor models: an overview of preclinical and translational research. Cancer Immun 13, 5 (2013).

23. W. L. Corwin et al., Tumor Control Index as a new tool to assess tumor growth in experimental animals. J Immunol Methods 445, 71–76 (2017).

24. J. Almeida-Santos et al., The multifaceted Foxp3(fgfp) allele enhances spontaneous and therapeutic immune surveillance of cancer in mice. Eur J Immunol 10.1002/eji.201948251 (2019).

25. K. Sakuishi et al., TIM3(+)FOXP3(+) regulatory T cells are tissue-specific promoters of T-cell dysfunction in cancer. Oncoimmunology 2, e23849 (2013).

26. J. C. Castle et al., Exploiting the mutanome for tumor vaccination. Cancer Res 72, 1081–1091 (2012).

27. G. M. Delgoffe et al., Stability and function of regulatory T cells is maintained by a neuropilin-1-semaphorin-4a axis. Nature 501, 252–256 (2013).

28. C. Wong, D. R. Goldstein, Impact of aging on antigen presentation cell function of dendritic cells. Curr Opin Immunol 25, 535–541 (2013).

29. D. Sauce, V. Appay, Altered thymic activity in early life: how does it affect the immune system in young adults? Curr Opin Immunol 23, 543–548 (2011).

30. T. E. Boursalian, J. Golob, D. M. Soper, C. J. Cooper, P. J. Fink, Continued maturation of thymic emigrants in the periphery. Nat Immunol 5, 418–425 (2004).

31. J. S. Hale, T. E. Boursalian, G. L. Turk, P. J. Fink, Thymic output in aged mice. Proc Natl Acad Sci U S A 103, 8447–8452 (2006).

32. G. Terszowski et al., Evidence for a functional second thymus in mice. Science 312, 284–287 (2006).

33. T. R. Simpson et al., Fc-dependent depletion of tumor-infiltrating regulatory T cells co-defines the efficacy of anti-CTLA-4 therapy against melanoma. J Exp Med 210, 1695–1710 (2013).

34. F. Arce Vargas et al., Fc Effector Function Contributes to the Activity of Human Anti-CTLA-4 Antibodies. Cancer Cell 33, 649–663 e644 (2018).

35. C. Raffin, L. T. Vo, J. A. Bluestone, Treg cell-based therapies: challenges and perspectives. Nat Rev Immunol 10.1038/s41577-019-0232-6 (2019).

36. M. L. Bergman et al., Tolerogenic insulin peptide therapy precipitates type 1 diabetes. J Exp Med 214, 2153–2156 (2017).

37. M. F. Moraes-Fontes et al., Steroid treatments in mice do not alter the number and function of regulatory T cells, but amplify cyclophosphamide-induced autoimmune disease. J Autoimmun 33, 109–120 (2009).

38. S. L. Silva et al., Autoimmunity and allergy control in adults submitted to complete thymectomy early in infancy. PLoS One 12, e0180385 (2017).

